# CONSENT: Scalable long read self-correction and assembly polishing with multiple sequence alignment

**DOI:** 10.1101/546630

**Authors:** Pierre Morisse, Camille Marchet, Antoine Limasset, Thierry Lecroq, Arnaud Lefebvre

## Abstract

**Motivation:** Third-generation sequencing technologies Pacific Biosciences and Oxford Nanopore allow the sequencing of long reads of tens of kbp, that are expected to solve various problems, such as contig and haplotype assembly, scaffolding, and structural variant calling. However, they also display high error rates that can reach 10 to 30%, for basic ONT and non-CCS PacBio reads. As a result, error correction is often the first step of projects dealing with long reads. As first long reads sequencing experiments produced reads displaying error rates higher than 15% on average, most methods relied on the complementary use of short reads data to perform correction, in a hybrid approach. However, these sequencing technologies evolve fast, and the error rate of the long reads now reaches 10 to 12%. As a result, self-correction is now frequently used as the first step of third-generation sequencing data analysis projects. As of today, efficient tools allowing to perform self-correction of the long reads are available, and recent observations suggest that avoiding the use of second-generation sequencing reads could bypass their inherent bias.

**Results:** We introduce CONSENT, a new method for the self-correction of long reads that combines different strategies from the state-of-the-art. More precisely, we combine a multiple sequence alignment strategy with the use of local de Bruijn graphs. Moreover, the multiple sequence alignment benefits from an efficient segmentation strategy based on *k*-mer chaining, which allows a considerable speed improvement. Our experiments show that CONSENT compares well to the latest state-of-the-art self-correction methods, and even outperforms them on real Oxford Nanopore datasets. In particular, they show that CONSENT is the only method able to efficiently scale to the correction of Oxford Nanopore ultra-long reads, and is able to process a full human dataset, containing reads reaching lengths up to 1.5 Mbp, in 15 days. Additionally, CONSENT also implements an assembly polishing feature, and is thus able to correct errors directly from raw long read assemblies. Our experiments show that CONSENT outperforms state-of-the-art polishing tools in terms of resource consumption, and provides comparable results. Moreover, we also show that, for a full human dataset, assembling the raw data and polishing the assembly afterwards is less time consuming than assembling the corrected reads, while providing better quality results.

**Availability and implementation:** CONSENT is implemented in C++, supported on Linux platforms and freely available at https://github.com/morispi/CONSENT.

## 1 Introduction

Third-generation sequencing technologies Pacific Biosciences (PacBio) and Oxford Nanopore Technologies (ONT) have become widely used since their inception in 2011. In contrast to second-generation technologies, producing reads reaching lengths of a few hundreds base pairs, they allow the sequencing of much longer reads (10 kbp on average [26], and up to >1 million bps [9]). These long reads are expected to solve various problems, such as contig and haplotype assembly [23, 10], scaffolding [3], and structural variant calling [27]. However, they are very noisy. More precisely, basic ONT and non-CCS PacBio reads can reach error rates of 10 to 30%, whereas second-generation short reads usually display error rates of 1%. The error profiles of these long reads are also much more complex than those of the short reads. Indeed, they are mainly composed of insertions and deletions, whereas short reads mostly contain substitutions. As a result, error correction is often required, as the first step of projects dealing with long reads. As the error profiles and error rates of the long reads are much different from those of the short reads, correcting long reads requires specific algorithmic developments.

The error correction of long reads has thus been tackled by two main approaches. The first approach, hybrid correction, makes use of additional short reads data to perform correction. The second approach, self-correction, aims at correcting the long reads solely based on the information contained in their sequences.

Hybrid correction methods rely on different techniques such as:

1. Alignment of short reads to the long reads (CoLoRMAP [8], HECiL [6])
2. Exploration of de Bruijn graphs, built from short reads *k*-mers (LoRDEC [24], Jabba [19], FMLRC [31])
3. Alignment of the long reads to contigs built from the short reads (MiRCA [11], HALC [1])
4. Hidden Markov Models, initialized from the long reads sequences and trained using the short reads (Hercules [7])
5. Combination of different strategies (NaS (1+3) [17], HG-CoLoR (1+2) [21])

Self-correction methods usually build around the alignment of the long reads against each other (PBDAGCon [5], PBcR [12]). We give further details on the state-of-the-art of self-correction in Section 1.1.

As first long reads sequencing experiments resulted in highly erroneous long reads (15-30% error rates on average), most methods relied on the additional use of short reads data. As a result, hybrid correction used to be much more widespread. Indeed, in 2014, for five hybrid correction tools, only two self-correction tools were available. However, third-generation sequencing technologies evolve fast, and now manage to produce long reads reaching error rates of 10-12%. Moreover, the evolution of long-read sequencing technologies also allows to produce higher throughputs of data, at a reduced cost. Consequently, such data became more widely available. As a result, self-correction is now frequently used as the first step of data analysis projects dealing with long reads.

### 1.1 Related works

Due to the fast evolution of third-generation sequencing technologies, and to the lower error rates they now reach, various efficient self-correction methods have recently been developed. Most of them share a common first step of computing overlaps between the long reads. This overlapping step can be performed in two different ways. First, a mapping approach can be used, to simply provide the positions of similar regions of the long reads (Canu [13], MECAT [32], FLAS [2]). Second, an alignment approach can be used, to provide the positions of similar regions, but also their actual base to base correspondence in terms of matches, mismatches, insertions, and deletions (PBDAGCon, PBcR, Daccord [29]). A directed acyclic graph (DAG) is then usually built, in order to summarize the 1V1 alignments and compute consensus, after recomputing actual alignments of mapped regions, if necessary. Other methods rely on de Bruijn graphs, either built from small windows of the alignments (Daccord), or directly from the long reads sequences with no prior overlapping step (LoRMA [25]). These graphs are explored, using the solid *k*-mers (*i.e. k*-mers occurring more frequently than a given threshold) from the reads as anchor points, in order to correct low quality, weak *k*-mers regions.

However, methods relying on direct alignment of the long reads are prohibitively time and memory consuming, and current implementations thus do not scale to large genomes. Methods solely relying on de Bruijn graphs, and avoiding the overlapping step altogether, usually require deep long reads coverage, since the graphs are usually built from large, solid *k*-mers. As a result, methods relying on a mapping strategy constitute the core of the current state-of-the-art for long read self-correction.

### 1.2 Contribution

We present CONSENT, a new self-correction method that combines different efficient approaches from the state-of-the-art. CONSENT indeed starts by computing multiple sequence alignments between overlapping regions of the long reads, in order to compute consensus sequences. These consensus sequences are then further polished with the help of local de Bruijn graphs, in order to correct remaining errors, and reduce the final error rate. Moreover, unlike current state-of-the-art methods, CONSENT computes actual multiple sequence alignments, using a method based on partial order graphs [14]. We also introduce an efficient segmentation strategy based on *k*-mer chaining, which allows to reduce the time footprint of the multiple sequence alignments. This segmentation strategy thus allows to compute scalable multiple sequence alignments. In particular, it allows CONSENT to efficiently scale to ONT ultra-long reads.

Our experiments show that CONSENT compares well to the latest state-of-the-art self-correction methods, and even outperforms them on real ONT datasets. In particular, they show that CONSENT is the only method able to efficiently scale to the correction of ONT ultra-long reads, and is able to perform correction on a full human dataset, containing reads reaching lengths up to 1.5 Mbp in 15 days.

Additionally, CONSENT is also able to polish assemblies generated from raw long reads. Our experiment on a full human dataset shows that assembling the raw data and polishing the assembly is less time consuming than assembling the corrected data, while offering better results. Our experiments also show that CONSENT outperforms state-of-the-art assembly polishing tools in terms of resource consumption, while providing comparable results.

## 2 Methods

### 2.1 Overview

CONSENT takes as input a FASTA file of long reads, and returns a set of corrected long reads, reporting corrected bases in uppercase, and uncorrected bases in lowercase. Like most efficient methods, CONSENT starts by computing overlaps between the long reads using a mapping approach. These overlaps are computed using an external program, and not by CONSENT itself. This way, only matched regions need to be further aligned in order to compute consensus. These matched regions are then divided into smaller windows, that are aligned independently. The alignment of these windows is performed via a multiple sequence alignment strategy based on partial order graphs. This multiple sequence alignment is computed by iteratively constructing and adding sequences to a DAG. It also benefits from an efficient heuristic, based on *k*-mer chaining, allowing to reduce the time footprint of computing multiple sequence alignments between noisy sequences. The DAG is then used to compute the consensus of the window it originates from. Once the consensus has been computed, a second step makes use of a local de Bruijn graph, in order to further polish it. This allows to correct weakly supported regions, that are, regions containing weak *k*-mers, and thus reduce the final error rate of the consensus. Finally, the consensus is realigned to the read, and correction is performed for each window. CONSENT’s workflow is summarized in Figure 1.

**Fig. 1:**
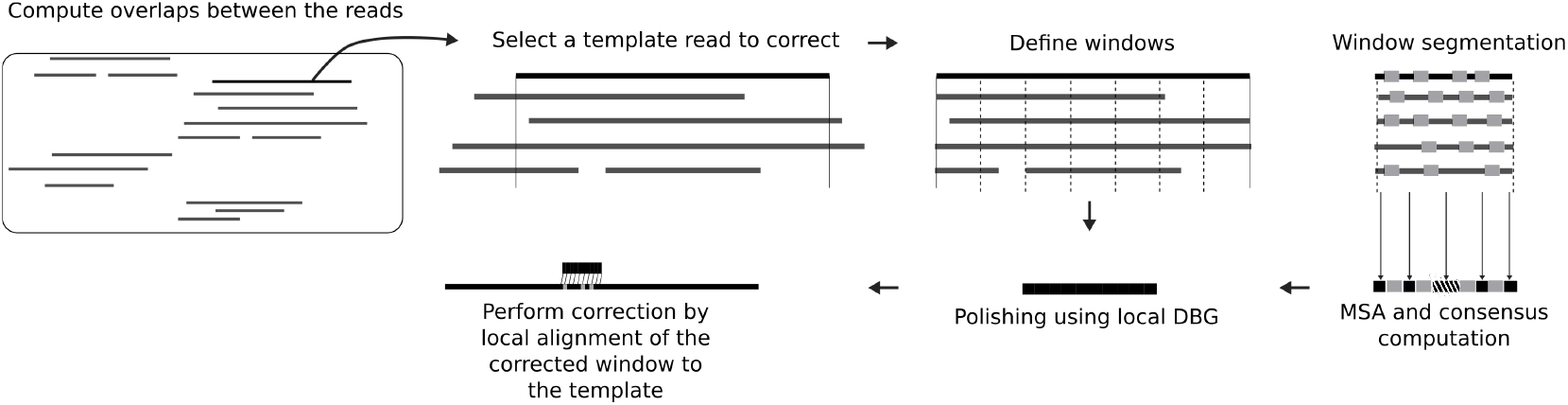
Overview of CONSENT’s workflow for long read error correction.

### 2.2 Definitions

Before presenting the CONSENT pipeline, we recall the notions of alignment piles and windows on such piles, as proposed in Daccord, since we rely on these throughout the rest of the paper.

#### 2.2.1 Alignment piles

An alignment pile represents a set of reads that overlap with a given read *A*. More formally, it can be defined as follows. For any given read *A*, we define an alignment pile for *A* as a set of alignment tuples (*A, R, Ab, Ae, Rb, Re, S*) where *R* is a long read id, *Ab* and *Ae* represent respectively the start and the end positions of the alignment on *A*, *Rb* and *Re* represent respectively the start and the end positions of the alignment on *R*, and *S* indicates whether *R* aligns forward (*S* = 0) or reverse complement (*S* = 1) to *A*. One can remark that this definition is slightly different from that of Daccord. In particular, Daccord adds an edit script to each tuple, representing the sequence of edit operations needed to transform *A*[*Ab..Ae*] into *R*[*Rb..Re*] if *S* = 0, or into 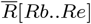 if *S* = 1 (where 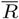 represents the reverse-complement of read *R*). This edit script can easily be retrieved by Daccord, as it relies on DALIGNER [22] to compute actual alignments between the long reads. However, as CONSENT relies on a mapping strategy, it does not have access to such information, and we thus chose to exclude the edit script from our definition of a tuple. In its alignment pile, we call the read *A* the *template* read. The alignment pile of a given template read *A* thus contains all the necessary information needed for its correction. An example of an alignment pile is given in Figure 2.

**Fig. 2:**
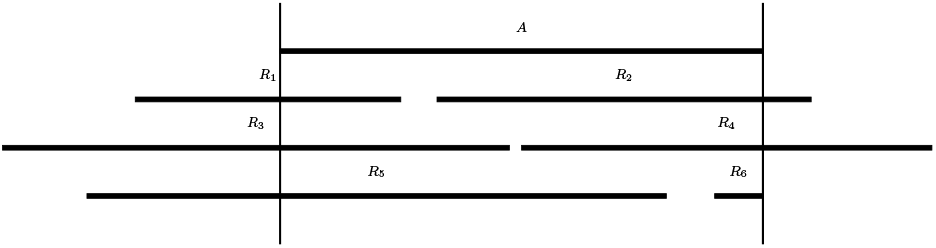
An alignment pile for a template read *A*. The pile is delimited by vertical lines at the extremities of *A*. Prefixes and suffixes of reads overlapping *A* outside of the pile are not considered during the next steps, as the data they contain will not be useful for correcting *A*.

### 2.2.2 Windows on alignment piles

In addition to the notion of alignment piles, Daccord also underlined the interest of processing windows from these piles instead of processing them as a whole. A window from an alignment pile is defined as follows. Given an alignment pile for a template read *A*, a window of this pile is a couple (*W_b_*, *W_e_*), where *W_b_* and *W_e_* represent respectively the start and the end positions of the window on *A*, and are such as 0 ≤ *W_b_* ≤ *W_e_* < |*A*| (*i.e.* the start and end positions of the window define a factor of the template read *A*). We refer to this factor as the *window’s template*. Additionally, in CONSENT, only windows having the two following properties are processed for correction:

- *W_e_* − *W_b_* + 1 = *L* (*i.e.* windows have a fixed size)
- ∀*i*, *W_b_* ≤ *i* ≤ *W_e_*, *A*[*i*] is supported by at least *C* reads of the pile, including *A* (*i.e.* windows have a minimum coverage threshold)

This second property allows to ensure that CONSENT has sufficient evidence to compute a reliable consensus for each window it processes. Examples of windows CONSENT does and does not process are given in Figure 3.

**Fig. 3:**
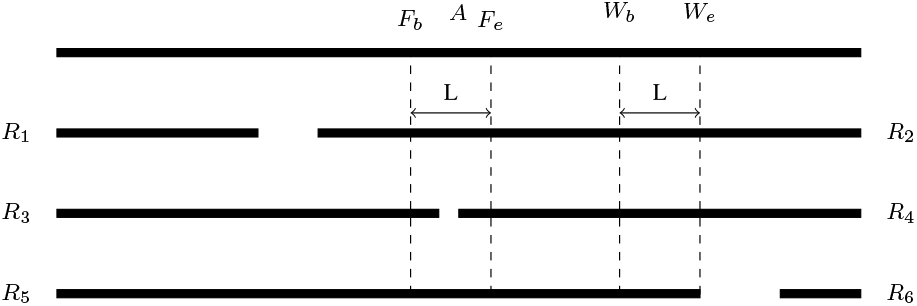
When fixing the length to *L* and the minimum coverage threshold to 4, the window (*W_b_*, *W_e_*) will be processed by CONSENT. With these same parameters, the window (*F_b_*, *F_e_*) will not be processed by CONSENT, as *A*[*i*] is not supported by at least 4 reads ∀ *F_b_* ≤ *i* ≤ *F_e_*.

In the case of Daccord, this window strategy allows to build local de Bruijn graphs with small values of *k*, and overcome the high error rates of the long reads, which cause issues when using large values of *k* [4]. More generally, processing windows instead of whole alignment piles allows to divide the correction problem into smaller subproblems that can be solved faster. Specifically, in our case, as we seek to correct long reads by computing multiple sequence alignments, working with windows allows to save both time and memory, since the sequences that need to be aligned are significantly shorter.

### 2.3 Overlapping

To avoid prohibitive computation time and memory consuming full alignments, CONSENT starts by overlapping the long reads using a mapping approach. By default, this step is performed with the help of Minimap2 [16]. However, CONSENT is not dependent on Minimap2, and the user can compute the overlaps with any other method, as long as the overlaps file follows the PAF format. We included Minimap2 as the default overlapper for CONSENT, since it offers good performances, and is thus able to scale to large organisms on reasonable setups.

### 2.4 Alignment piles and windows computation

The alignment piles are computed by parsing the PAF file generated by the overlapper during the previous step. Each line indeed contains all the necessary information to define a tuple from an alignment pile. It includes the identifiers of the two long reads, the start and the end positions of their overlap, as well as the orientation of the second read relatively to the first. Moreover, for each alignment pile, CONSENT only includes the *N* highest identity overlaps (*N* = 150 by default, although it can be user-specified), in order to reduce the time footprint, and avoid computing costly multiple sequence alignments of numerous sequences.

Given an alignment pile for a read *A*, we can then compute its set of windows. To this aim, we use an array of length *|A|*, which counts how many times each nucleotide of *A* is supported. We initialize the array with 1s at each position, and for each tuple (*A, R, Ab, Ae, Rb, Re, S*), we increment values at positions *i* such as *Ab* ≤ *i* ≤ *Ae*. After processing all the tuples, we retrieve the positions of the piles by finding, in the array, sketches of length *L* of values ≥ *C*. We search for such sketches because CONSENT only processes windows of fixed length and with a minimum coverage threshold. In practice, we extract overlapping windows instead of partitioning the pile into a set of non-overlapping windows. Indeed, since it is usually harder to exploit alignments located on sequences extremities, consensus sequence might be missing at the extremities of some windows. Such events would thus cause a lack of correction on the reads, and using overlapping windows allows to overcome the issue. Each window is then processed independently during the next steps. Moreover, the reads are loaded into memory to support random access and thus accelerate the correction process. Each base is encoded using 2 bits in order to reduce memory usage. The memory consumption is thus roughly 1/4 of the total size of the reads.

### 2.5 Window consensus

We process each window in two distinct steps. First, we align the sequences from the window using a multiple sequence alignment strategy based on partial order graphs, in order to compute consensus. This multiple sequence alignment strategy also benefits from an efficient heuristic, based on *k*-mer chaining, allowing to decompose the global problem into smaller instances, thus reducing both time and memory consumption. Second, after computing the window’s consensus, we further polish it with the help of a local de Bruijn graph, at the scale of the window, in order to get rid of the few errors that might remain despite consensus computation.

#### 2.5.1 Consensus computation

In order to compute the consensus of a window, CONSENT uses POAv2 [14], an implementation of a multiple sequence alignment strategy based on partial order graphs. These directed acyclic graphs, store all the information of the multiple sequence alignment. This way, at each step (*i.e.* at each alignment of a new sequence), the graph contains the current multiple sequence alignment result. To add a new sequence to the multiple sequence alignment, the sequence is aligned to the DAG, using a generalization of the Smith-Waterman algorithm.

Other methods usually compute 1V1 alignments between the read to be corrected and other reads overlapping with it, and then build a result DAG to summarize the alignments, and represent the multiple sequence alignment. In contrast, CONSENT’s strategy allows us to compute actual multiple sequence alignments, and to directly build the DAG, during the alignment computation. Indeed, the DAG is first initialized with the sequence of the window’s template, and is then iteratively enriched by aligning the other sequences from the window, until it becomes the final, result graph. We then extract a matrix, representing the multiple sequence alignment, from the graph, and compute consensus by performing a majority vote. When a tie occurs, we chose the nucleotide from the window’s template as the consensus base.

However, even on small windows, computing multiple sequence alignments on hundreds of bases from dozens of sequences is computationally expensive, especially when the divergence among sequences is high. To avoid the burden of building a consensus by computing full multiple sequence alignments, we search for collinear regions shared by these sequences, in order to split the global task into several smaller instances. We thus build several consensus on regions delimited by anchors shared among the sequences, and reconstruct the global consensus from the distinct, smaller consensus sequences obtained. The rationale is to benefit from the knowledge that all the sequences come from the same genomic area. This way, on the one hand, we can compute multiple sequence alignments of shorter sequences, which greatly reduces the computational costs. On the other hand, we only use related sequences to build the consensus, and therefore exclude spurious sequences. This behavior allows a massive speedup along with an improvement in the global consensus quality.

To find such collinear regions, we first select *k*-mers that are non-repeated in their respective sequences, and shared by multiple sequences. We then rely on dynamic programming to compute the longest anchors chain *a*_1_, …, *a_n_* such as:

1. ∀*i, j* such that 1 ≤ *i* < *j* ≤ *n*, *a_i_* appears before *a_j_* in every sequence containing *a_i_* and *a_j_*
2. ∀*i*, 1 ≤ *i* < *n*, there are at least *T* reads containing *a_i_* and *a*_*i*+1_ (with *T* a solidity threshold equal to 8 by default).

We therefore compute multiple, local consensus, using substrings bordered by consecutive anchors, in sequences that contain them, and are then able to reconstruct the global consensus of the window: *consensus*(*prefix*) + *a_i_* + *consensus*(]*a*_1_, *a*_2_[) + *a*_2_ + · · · + *consensus*(]*a*_*n*−1_, *a*_*n*_[)+*a*_*n*_+*consensus*(*suffix*). We illustrate this segmentation strategy in Supplementary Figures S1 (longest anchors chain computation) and S2 (local consensus computation and global consensus reconstruction).

#### 2.5.2 Consensus polishing

After processing a given window, a few erroneous bases might remain on the computed consensus. This might happen in cases where the coverage depth of the window is relatively low, and thus cannot yield a high-quality consensus. Consequently, we propose an additional, second correction phase, that aims at polishing the consensus obtained during the previous step. This allows CONSENT to further enhance its quality, by correcting weakly supported *k*-mers. This feature is related to Daccord’s local de Bruijn graph correction strategy.

First, a local de Bruijn graph is built from the window’s sequences, using only small, solid, *k*-mers. The rationale is that small *k*-mers allows CONSENT to overcome the classical issues encountered due to the high error rate of the long reads, when using large *k* values. CONSENT then searches for regions only composed of weak *k*-mers, flanked by sketches of *n* (usually, *n* = 3) solid *k*-mers. Afterwards, CONSENT attempts to find a path allowing to link a solid *k*-mer from the left flanking region to a solid *k*-mer from the right flanking region. We call these solid *k*-mers *anchors*. The graph is thus traversed, in order to find a path between two anchors, using backtracking if necessary. If a path between two anchors is found, the region containing the weak *k*-mers is replaced by the sequence dictated by this path. If none of the anchors pairs can be linked, the region is left unpolished. To polish sketches of weak *k*-mers located at the left (respectively right) extremity of the consensus, highest weighted edges of the graph are followed, until the length of the path reaches the length of the region to polish, or no edge can be followed out of the current node.

### 2.6 Read correction via window consensus alignment

Once the consensus of a window has been computed and polished, we need to realign it to the template, in order to actually perform correction. To this aim, we use an optimized library of the Smith-Waterman algorithm [33]. To avoid time-costly alignment, we locally align the consensus around the positions of the window it originates from. This way, given a window (*W_b_*, *W_e_*) of the alignment pile of the read *A*, its consensus will be aligned to *A*[*W_b_* − *O*..*W_e_* + *O*], where *O* represents the length of the overlap between consecutive windows processed by CONSENT (*O* = 50 by default, although it can be user-specified). Aligning the consensus outside of the original window’s extremities as such allows to take into account the error profile of the long reads. Indeed, as insertions and deletions are predominant in long reads, it is likely that a consensus could be longer than the window it originates from, thus spanning outside of this window’s extremities.

In the case alignment positions of the consensus from the *i*th window overlap with alignment positions of the consensus from the (*i* + 1)th window, we compute the overlapping sequences of the two consensus. The one containing the largest number of solid *k*-mers (where the *k*-mer frequencies of each sequence are computed from the window their consensus originate from) is chosen and kept as the correction. In the case of a tie, we arbitrarily chose the sequence from the (*i* + 1)th consensus as the correction. We then correct the aligned factor of the long read by replacing it with the aligned factor of the consensus.

## 3 Experimental results

### 3.1 Impact of the segmentation strategy

Before comparing CONSENT to state-of-the-art self-correction tools, we first validate our segmentation strategy. To this aim, we simulated a 50x coverage of long reads from *E.coli*, with a 12% error rate, using SimLoRD [28]. The following parameters were used for the simulation: `–probability-threshold 0.3 −prob-ins 0.145 −prob-del 0.06, and −prob-sub 0.02`. We then ran the CONSENT pipeline, with, and without the segmentation strategy. Results of this experiment are given in Table 1. We obtained these results using ELECTOR [18], a tool specifically designed to precisely measure correction accuracy on simulated data. In particular, ELECTOR outputs metrics such as recall, precision, and error rate before and after correction. These results show that, in addition to being 47x faster than the regular MSA implementation, our segmentation strategy also allows us to reach slightly lower memory consumption, as well as higher quality. In particular, the post-correction error rate is divided by 1.77, and the precision increases by almost 0.15% when using the segmentation strategy. This gain in quality can be explained by the fact that our segmentation strategy allows to get rid of spurious sequence and thus to compute more accurate alignments and consensus.

**Table 1.**
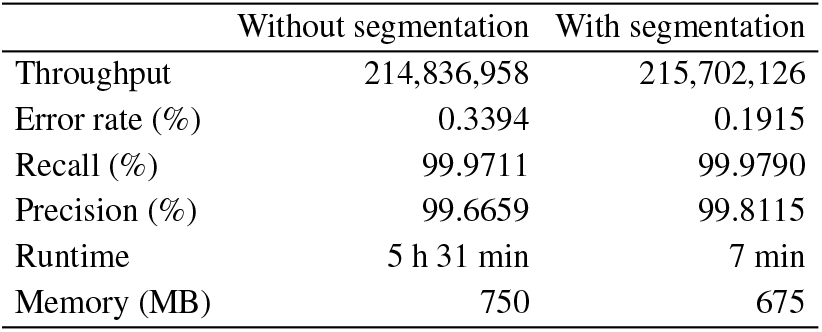
Comparison of the results produced by CONSENT, with and without our segmentation stategy, as reported by ELECTOR. Using the segmentation strategy allows a 47x speed-up, while producing a slightly higher quality correction.

|  | Without segmentation | With segmentation |
| --- | --- | --- |
| Throughput | 214,836,958 | 215,702,126 |
| Error rate (%) | 0.3394 | 0.1915 |
| Recall (%) | 99.9711 | 99.9790 |
| Precision (%) | 99.6659 | 99.8115 |
| Runtime | 5 h 31 min | 7 min |
| Memory (MB) | 750 | 675 |

### 3.2 Comparison to the state-of-the-art

We now compare CONSENT against state-of-the-art error correction methods. We include the following tools in the benchmark: Canu, Daccord, FLAS, and MECAT. We voluntarily exclude LoRMA from the comparison, as it tends to aggressively split the reads, and thus produce reads that are usually shorter than 900 bp. We however report LoRMA’s result in Supplementary Tables S3 and S4. We also exclude hybrid error correction tools from the benchmark, as we believe it makes more sense to only compare self-correction tools. We performed experiments both on simulated and real data. Comparison on simulated data is presented in Section 3.2.2, and comparison on real data in Section 3.2.3. Datasets used for the different experiments are presented in Section 3.2.1. We ran all tools with default or recommended parameters. For CONSENT, we set the minimum support to define a window to 4, the window size to 500, the overlap size between consecutive windows to 50, the *k*-mer size used for chaining and polishing to 9, the solidity threshold for *k*-mers to 4, and the solidity threshold for the anchors chain computation to 8. Additionally, only windows for which at least two anchors could be found during the segmentation algorithm were processed.

#### 3.2.1 Datasets

For our experiments, we used both simulated PacBio and real ONT long reads. PacBio reads were simulated with SimLoRD, using the same parameters as in Section 3.1. We generated two datasets with a 12% error rate for *E. coli*, *S. cerevisiae* and *C. elegans*: one with a 30x coverage, and one with a 60x coverage, corresponding to typical sequencing depths in current long reads experiments. As for the real ONT data, we used a 63x coverage dataset from *D. melanogaster*, and a 29x coverage from *H. sapiens* chr 1, containing ultra-long reads, reaching lengths up to 340 kbp. Further details and accession numbers for all the datasets are given in Supplementary Table S1. Details on the reference sequences are given in Supplementary Table S2.

#### 3.2.2 Comparison on simulated data

To precisely assess the accuracy of the different correction methods, we first tested them on the simulated PacBio datasets. ELECTOR was used to evaluate the correction accuracy of each method. Correction statistics of all the aforementioned tools on the different datasets, along with their runtime and memory consumption, are given in Table 2. For methods having distinct, easily identifiable, steps for overlapping and correction (*i.e.* Daccord, MECAT and CONSENT), we additionally report runtime and memory consumption of these two processes apart. We ran all the correction experiments on a computer equipped with 16 2.39 GHz cores and 32 GB of RAM.

**Table 2.**
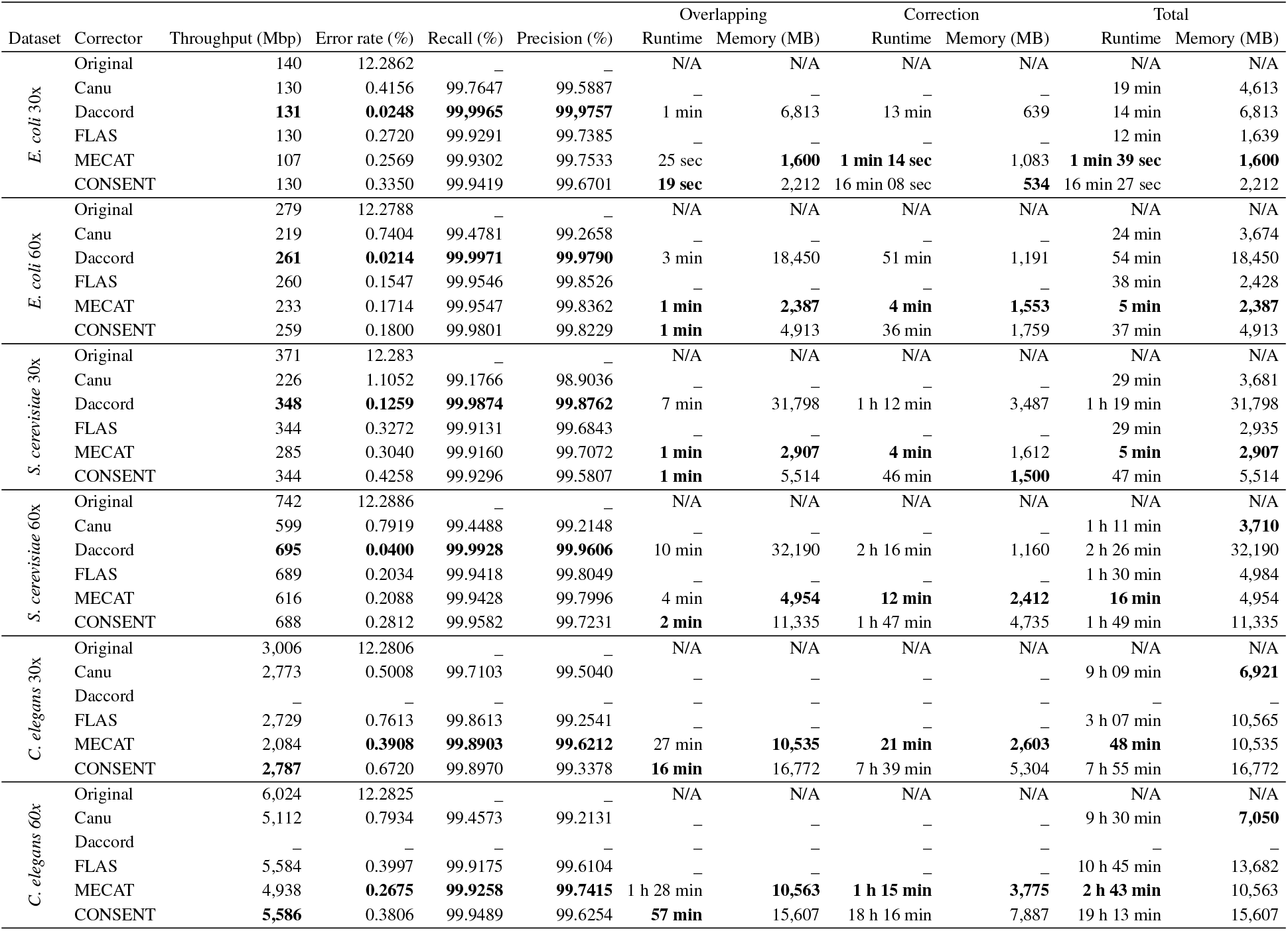
Metrics output by ELECTOR on the simulated PacBio datasets. Daccord results are missing for the two C. elegans datasets, as DALIGNER failed to perform alignment, reporting an error upon start, even when ran on a cluster node with 28 2.4 GHz cores and 128 GB of RAM. Recall and precision are not reported for original reads, since they cannot be computed from uncorrected reads.

Daccord clearly performed the best in terms of throughput and quality, outperforming all the other methods on the *E. coli* and the *S. cerevisiae* datasets. However, the overlapping step, relying on actual alignment of the long reads against each other, consumed high amounts of memory, 3x to 11x more than CONSENT or MECAT mapping strategies. As a result, Daccord could not scale to the *C. elegans* datasets, DALIGNER reporting an error upon start, even when run on a cluster node equipped with 128 GB of RAM. On the contrary, Canu displayed the highest error rates on all the datasets, except on the *C. elegans* dataset with a 30x coverage, but consumed relatively stable, low amounts of memory. In particular, on the two *C. elegans* datasets, it displayed the lowest memory consumption among all the other methods.

MECAT performed the best in terms of runtime, outperforming all the other tools on all the datasets. Its overlapping strategy was also highly efficient, and displayed the lowest memory consumption among all the other strategies, on all the datasets. However, compared to Minimap2 (the overlapping strategy adopted in CONSENT) MECAT’s overlapping strategy displayed higher runtimes, although it remained faster than Daccord’s DALIGNER. Minimap2’s memory consumption, however, was larger than that of MECAT’s overlapping strategy, on all the datasets. The memory consumption of Minimap2 can nonetheless easily be reduced, at the expense of a slightly larger runtime, by decreasing the size of the index used for computing the overlaps, which CONSENT sets to 1 Gbp by default.

Compared to both FLAS and CONSENT, MECAT displayed lower throughputs on all the datasets. As for FLAS, this can be explained by the fact that it is a MECAT wrapper, allowing to retrieve additional overlaps, and thus correct a greater number long reads. As a result, since it relies on MECAT’s error correction strategy, FLAS displayed highly similar memory consumption. Runtime was however higher, due to the additional steps allowing to retrieve supplementary overlaps, and to the resulting higher number of reads to correct. Throughputs and error rates of FLAS and CONSENT were highly similar on all the datasets, varying by 0.1% at most, on the *S. cerevisiae* dataset with a 30x coverage. Runtimes were also comparable on the *E. coli* and *S. cerevisiae* datasets. However, on the *C. elegans* datasets, CONSENT displayed higher runtimes. As for the memory consumption of the error correction step, CONSENT was less efficient than MECAT on most datasets. This can be explained by the fact that CONSENT loads the correction jobs into a thread pool of default size 100,000. Reducing the size of the thread pool would allow CONSENT to consume less memory, at the expense of a slightly higher runtime.

#### 3.2.3 Comparison on real data

We then evaluated the different correction methods on larger, real ONT datasets. For these datasets, we not only evaluate how well the corrected long reads realign to the reference genome, but also how well they assemble. For the alignment assessment, we report how many reads were corrected, their throughput, their N50, the proportion of corrected reads that could be aligned, the average identity of the alignments, as well as the genome coverage, that is, the percentage of bases of the reference genome to which at least a nucleotide aligned. For the assembly assessment, we report the overall number of contigs, the number of contigs that could be aligned, the NGA50 and NGA75, and, once again, the genome coverage. We obtained alignment statistics using ELECTOR’s second module, which performs alignment to the reference genome with Minimap2. We performed assemblies using Minimap2 and Miniasm [15], and obtained statistics with QUAST-LG [20]. Results are given in Table 3 for the alignment assessment, and in Table 4 for the assembly assessment. Runtimes and memory consumption of the different methods are also given in Table 3. As for the simulated data, we report runtime and memory consumption of the overlapping and correction steps apart, when possible. We ran all the correction experiments on a cluster node equipped with 28 2.39 GHz cores and 128 GB of RAM.

**Table 3.**
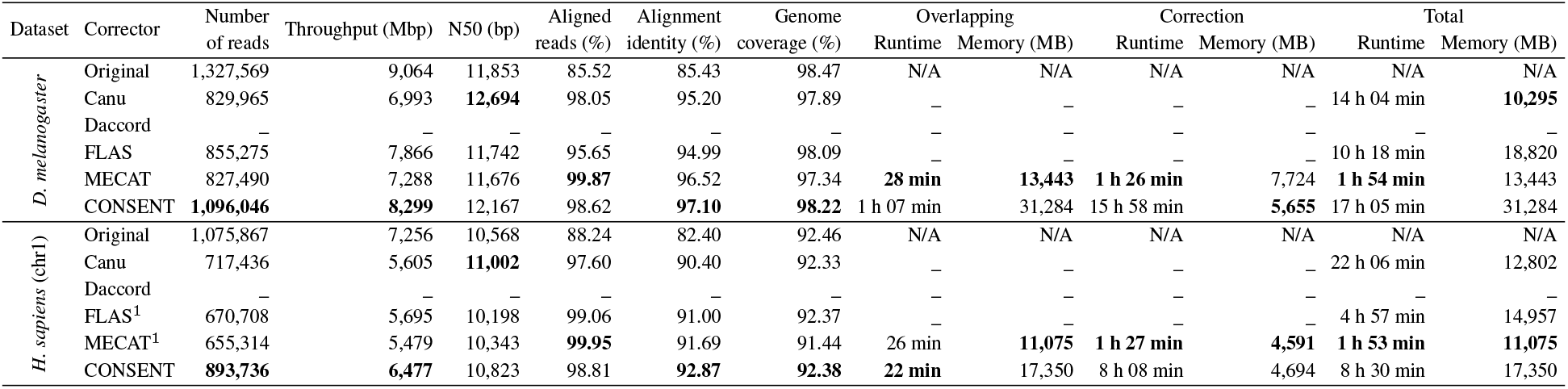
Statistics of the real long reads, before and after correction with the different methods. ^1^ Reads longer than 50 kbp were filtered out, as ultra-long reads caused the programs to stop with an error. There were 1,824 such reads in the original dataset, accounting for a total number of 135,364,312 bp. Daccord could not be run on these two datasets, due to errors reported by DALIGNER.

**Table 4.** Statistics of the assemblies generated from the raw and corrected long reads. As previously mentioned, Daccord results on the two datasets are absent, since it could not be run. For the assembly of the original reads on the H. sapiens (chr 1) dataset, QUAST-LG did not provide a metric for the NGA75.

| Dataset | Corrector | Contigs | Aligned contigs (%) | NGA50 (bp) | NGA75 (bp) | Genome coverage (%) | Errors / 100 kbp |
| --- | --- | --- | --- | --- | --- | --- | --- |
| *D. melanogaster* | Original | 423 | 96.45 | 864,011 | 159,590 | 83.22 | 10,690 |
|  | Canu | 410 | 92.93 | 2,757,690 | 822,577 | **92.95** | 1,896 |
|  | Daccord | – | – | – | – | – | – |
|  | FLAS | 407 | 98.53 | 1,123,346 | 363,017 | 92.16 | 2,736 |
|  | MECAT | **310** | **99.68** | 1,414,076 | 480,297 | 92.02 | 1,731 |
|  | CONSENT | 348 | 98.56 | **4,081,831** | **958,935** | 92.06 | **1,549** |
| *H. sapiens* (chr1) | Original | 201 | 93.53 | 1,008,692 | – | 77.52 | 11,318 |
|  | Canu | 361 | 98.61 | 946,029 | 245,015 | 94.85 | 4,689 |
|  | Daccord | – | – | – | – | – | – |
|  | FLAS | 237 | 100.00 | 1,698,601 | 289,968 | 94.97 | 4,404 |
|  | MECAT | 259 | **100** | 1,378,242 | 287,113 | **94.89** | **3,413** |
|  | CONSENT | **166** | 93.37 | **2,385,471** | **580,981** | 91.76 | 4,669 |

On these two datasets, Daccord failed to run, as DALIGNER could not perform alignment, for the same reason as for the simulated *C. elegans* datasets. CONSENT corrected the largest number of reads, and reached the highest alignment identity on the two datasets. Its N50 was also higher than that of all the other methods, except Canu. CONSENT also reached the highest throughput, and the largest genome coverage, for the two datasets. When it comes to runtime and memory consumption, MECAT once again outperformed all the other methods, as in the experiments on simulated data. Moreover, it reached the highest proportion of aligned reads, on both datasets. However, CONSENT was close, since only 1.14-1.25% fewer reads could be aligned.

Moreover, on the *H. sapiens* (chr 1) dataset, CONSENT and Canu were the only tools able to deal with ultra-long reads. Indeed, other methods reported errors when attempting to correct the original dataset. As a result, in order to allow these methods to perform correction, we had to manually remove the reads longer than 50 kbp. There were 1,824 such reads, accounting for a total number of 135,364,312 bp. However, even if it managed to scale to the correction of ultra-long reads, Canu was almost four times slower than CONSENT, making CONSENT the only tool to efficiently scale to ultra-long reads.

On the *D. melanogaster* dataset, the assembly yielded from Canu corrected reads slightly outperformed all the other assemblies in terms of genome coverage. However, it was composed of a higher number of contigs compared to all the other assemblies, except the one obtained from the raw reads. The assembly obtained from CONSENT corrected reads outperformed all the other assemblies in terms of NGA50, NA75 as well as error rate per 100 kbp. The genome coverage of the CONSENT assembly was also slightly larger than that of FLAS and MECAT.

On the *H. sapiens* (chr 1) dataset, the assembly obtained from CONSENT corrected reads outperformed all the other assemblies in terms of number of contigs, NGA50, and NGA75. In particular, the NGA50 of the CONSENT assembly was almost 700 kbp larger than that of other assemblies. However, 11 contigs of the CONSENT assembly could not be aligned to the reference. As a result, compared to the assemblies obtained from FLAS and MECAT corrected reads, the assembly yielded from the CONSENT corrected reads covered 4% less of the reference sequence, and displayed a higher error rate per 100 kbp. These unaligned contigs and differences could likely be reduced by further adapting both CONSENT and Miniasm parameters.

### 3.3 Assembly polishing

As an additional feature, CONSENT also allows to perform assembly polishing. The process is pretty straightforward. Indeed, instead of computing overlaps between the long reads, as presented in the previous sections, overlaps are simply computed between the assembled contigs and the long reads used for the assembly. The rest of the pipeline remains the same. We present assembly polishing results on the simulated *E. coli*, *S. cerevisiae*, and *C. elegans* datasets with a 60x coverage, as well as on the real *D. melanogaster* and *H. sapiens* (chr 1) datasets. We compare CONSENT to RACON [30], a state-of-the-art assembly polishing method. Results are presented in Table 5.

**Table 5.** Statistics of the assemblies, before and after polishing with RACON and CONSENT. The missing contig for the CONSENT and RACON polishings on the D. melanogaster dataset is 428 bp long, and could not be polished, due to the window size of the two methods being larger (500).

| Dataset | Method | Contigs | Aligned contigs (%) | NGA50 (bp) | NGA75 (bp) | Genome coverage (%) | Errors / 100 kbp | Runtime | Memory (MB) |
| --- | --- | --- | --- | --- | --- | --- | --- | --- | --- |
| *E. coli* 60x | Original | **1** | **100.00** | 4,939,014 | 4,939,014 | **99.91** | 10,721 | N/A | N/A |
|  | RACON | **1** | **100.00** | 4,663,914 | 4,663,914 | 99.90 | 499 | 5 min 55 sec | **643** |
|  | CONSENT | **1** | **100.00** | 4,637,850 | 4,637,850 | **99.91** | **117** | **33 sec** | 971 |
| *S. cerevisiae* 60x | Original | **29** | **100.00** | 579,247 | 456,470 | **96.14** | 10,694 | N/A | N/A |
|  | RACON | **29** | **100.00** | 539,472 | 346,116 | 96.09 | 637 | 15 min 47 sec | 1,703 |
|  | CONSENT | **29** | **100.00** | 532,246 | 332,862 | 96.07 | **208** | **3 min 30** | **1,336** |
| *C. elegans* 60x | Original | **47** | **100.00** | 5,201,998 | 2,511,520 | **99.78** | 10,974 | N/A | N/A |
|  | RACON | **47** | 97.87 | 6,405,523 | 2,726,529 | 99.74 | 819 | 2 h 24 min | 14,288 |
|  | CONSENT | **47** | 97.87 | 6,336,755 | 2,699,456 | 99.72 | **410** | **14 min** | **4,379** |
| *D. melanogaster* | Original | **423** | 96.45 | 864,011 | 159,590 | 83.20 | 10,690 | N/A | N/A |
|  | RACON | 422 | 98.34 | **1,446,703** | **552,532** | **93.03** | **961** | 3 h 29 min | 19,508 |
|  | CONSENT | 422 | **98.82** | 1,235,674 | 449,364 | 91.94 | 2,240 | **1 h 51 min** | **5,814** |
| *H. sapiens* (chr 1) | Original | **201** | 93.53 | 1,008,692 | — | 77.52 | 11,318 | N/A | N/A |
|  | RACON | **201** | 97.01 | **3,481,900** | **1,282,763** | **95.69** | **2,393** | 2 h 30 min | 16,202 |
|  | CONSENT | **201** | **97.57** | 3,283,830 | 891,052 | 93.88 | 4,805 | **59 min** | **7,101** |

These results show that CONSENT outperformed RACON in terms of quality of the results, especially dealing better with errors, and thus greatly reducing the error rate per 100 kbp, on the *E. coli*, *S. cerevisiae*, and *C. elegans* datasets. Moreover, the NGA50, NGA75 and genome coverage of CONSENT were highly similar to those of RACON on these three datasets.

For the larger, eukaryotic *D. melanogaster* dataset, RACON outperformed CONSENT in terms of error rate and genome coverage, but the NGA50, NGA75 of the two methods remained comparable. On the *H. sapiens* (chr 1) dataset, RACON once again outperformed CONSENT in terms of error rate and genome coverage, and also displayed larger NGA50 and NGA75. However, polishing the assembly with CONSENT allowed to align a greater proportion of contigs, compared to both the raw and the RACON polished assembly. Additionally, on all the datasets, CONSENT was 2x to 11x faster than RACON, and also consumed up to three times less memory.

### 3.4 Results on a full human dataset

To further validate the scalability of CONSENT, we present results on a full ONT human dataset. This dataset is composed of 113 Gbp, displays an error rate of 17%, and contains ultra-long reads reaching lengths up to 1.5 Mbp. Further details are given in Supplementary Table S1. In this experiment, we not only evaluate how CONSENT behaves on such a large dataset, but also study the impact of the correction / assembly order on the quality of the results. We thus correct the raw data with CONSENT, and then assemble the corrected long reads, but also assemble the raw long reads first, and then polish the assembly with CONSENT. Alignment statistics of the raw and corrected long reads are presented in Table 6, while statistics of the different assemblies are presented in Table 7.

**Table 6.** Statistics of the full H. sapiens dataset, before and after correction with CONSENT.

| Corrector | Number of reads | Throughput (Mbp) | N50 (bp) | Aligned reads (%) | Alignment identity (%) | Genome coverage (%) | Overlapping Runtime | Overlapping Memory (MB) | Correction Runtime | Correction Memory (MB) | Total Runtime | Total Memory (MB) |
| --- | --- | --- | --- | --- | --- | --- | --- | --- | --- | --- | --- | --- |
| Original | 15,243,243 | 112,970 | 12,196 | 80.57 | 82.74 | 93.56 | N/A | N/A | N/A | N/A | N/A | N/A |
| CONSENT | 11,913,704 | 102,165 | 12,826 | 98.43 | 93.66 | 93.33 | 8 days 11 h | 98,324 | 6 days 15 h | 46,397 | 15 days 2 h | 98,324 |

**Table 7.** Statistics of the different assemblies for the full H. sapiens dataset. Raw corresponds to the assembly generated from raw reads. Corrected corresponds to the assembly generated from corrected reads. Polished corresponds to the assembly generated from raw reads, and polished with CONSENT. Runtime and memory consumption are reported for the whole correction + assembly or assembly + polishing pipelines. QUAST-LG did not provide a metric for the NGA75 of the assembly generated from corrected reads.

| Assembly | Contigs | Aligned contigs (%) | NGA50 (bp) | NGA75 (bp) | Genome coverage (%) | Errors / 100 kbp | Runtime | Memory (MB) |
| --- | --- | --- | --- | --- | --- | --- | --- | --- |
| Raw | 750 | 95.47 | 534,347 | — | 69.83 | 11,175 | 2 days 20 h | 382,191 |
| Corrected | 911 | 97.69 | 1,837,315 | — | 77.36 | 5,862 | 22 days 2 h | 1,058,085 |
| Polished | 749 | 98.13 | 2,869,858 | 176,799 | 83.63 | 4,654 | 7 days 15 h | 382,191 |

Alignment statistics of Table 6 show that CONSENT managed to process the whole dataset in 15 days, and required less than 100 GB of RAM. More precisely, the more computationally expensive step, both in terms of runtime and memory consumption, was actually the overlaps computation, and not the error correction itself. The corrected reads displayed a higher N50 than the raw reads, the longest read reaching 929 kbp, and almost 99% of them could be realigned to the reference genome. The average identity of the alignments reached more than 93.5%, which is slightly higher, but consistent with the results on chr 1, presented in Table 3. Moreover, CONSENT managed to correct a large number of reads, and thus barely reduced the genome coverage of the original dataset.

Assemblies statics of Table 7 are particularly interesting. Indeed, they show that, in addition to being extremely more computationally expensive, correcting the reads before assembling them produces less satifying results than assembling the raw reads first, and then polishing the assembly. Indeed, the correction + assembly pipeline required more than 22 days and 1 TB of RAM, while the assembly + polishing pipeline ran in less than 8 days, and consumed less than 400 GB of RAM. In addition, the polished assembly displayed better metrics than the assembly generated from corrected reads, reaching higher NGA50, NGA75, and genome coverage, and lower error rate per 100 kbp. These results underline the fact that, for large datasets and complex genomes, assembling the raw data first, and then polishing the assembly is much more efficient than correcting the reads and then performing assembly.

## 4 Discussion and future works

Experimental results on the human datasets are particularly promising. Indeed, they show that CONSENT is the only method able to efficiently scale to the ultra-long reads they contain. More precisely, on the human chr 1 dataset, CONSENT is almost four times faster than Canu, the only other method able to scale to the correction of ultra-long reads. Moreover, it also produces more accurate results, and thus allows to yield a more contiguous assembly. As such reads are expected to become more widely available in the future, being able to deal with them will soon become a necessity. In addition, results on the complete human dataset show that CONSENT manages to efficiently process such large datasets in 15 days, using less than 100 GB of RAM. Moreover, this memory consumption could easily be reduced by adapting the parameters of Minimap2, and reducing the size of the thread pool used during the actual correction step. At the expense of an increased runtime, CONSENT could thus process a full human dataset on a simple laptop. Further experiments should therefore focus on the correction of larger and more complex organisms. However, the runtime of CONSENT’s correction step tends to be higher than that of other state-of-the-art methods. We discuss how to further reduce these computational costs below.

Our experiments show that the runtime of the correction step tends to rise according to the complexity of the genome. This can be explained by the highest proportion of repeated regions in more complex genomes. Such repeated regions indeed impact the alignment piles coverages, and could therefore lead to the processing of piles having very deep coverages. For such piles, our strategy of only selecting the *N* highest identity overlaps might prove inefficient, especially when the length of the repeated regions grows longer. To further refine the overlaps selection, we could use a validation strategy similar to that of HALC. Such a strategy would allow us to only consider sequences from the pile that actually come from the same genomic region as the long read we are attempting to correct. This would, in turn, allow us to ensure the selected sequences display low divergence, which would speed up the multiple sequence alignment computation, while allowing to produce higher quality consensus.

Moreover, further optimization of the parameters shall also be considered. In particular, the window size and the minimum number of anchors to allow the processing of a window significantly impact the runtime. Running various experiments with different sets of parameters could therefore allow us to find a satisfying compromise between runtime and quality of the results. The fact that the CONSENT assembly covers a smaller proportion of the reference sequence also gives us further room for improvement. In particular, looking to the unaligned contigs more into details could help us further improve the mechanisms and principles of CONSENT. Another possible improvement would be to consider multiple *k*-mer size for the *k*-mer chaining strategy. By selecting the best possible chaining according to the coverage or the repetitive elements of a given window, the method could be more robust and more efficient by computing smaller multiple sequence alignments.

Finally, it is essential to note that, as mentioned in Section 2.3, CONSENT uses Minimap2 as its default overlapper, but does not depend on this tool. As a result, CONSENT will benefit from the progress of future overlapping strategies, and will therefore allow to propose better correction quality as the overlapping methods evolve.

## 5 Conclusion

We presented CONSENT, a new self-correction method for long reads that combines different efficient strategies from the state-of-the-art. CONSENT starts by computing overlaps between the long reads to correct. It then divides the overlapping regions into smaller windows, in order to compute multiple sequence alignments, and consensus sequences of each window independently. These multiple sequence alignments are performed using a method based on partial order graphs, allowing to perform actual multiple sequence alignment. This method is combined to an efficient *k*-mer chaining strategy, which allows to further divide the multiple sequence alignments into smaller instances, and thus significantly reduce computation times. After computing the consensus of a given window, it is further polished with the help of a local de Bruijn graph, at the scale of the window, in order to further reduce the final error rate. Finally, the polished consensus is locally realigned to the read, in order to correct it.

Our experiments show that CONSENT compares well to, or even outperforms, other state-of-the-art self-correction methods in terms of quality of the results. In particular, CONSENT is the only method able to efficiently scale to the correction of ONT ultra-long reads, and is able to process a full human dataset containing reads reaching lengths up to 1.5 Mbp in 15 days. Although very recent, such reads are expected to further develop, and thus become more widely available in the near future. Being able to deal with them will thus soon become a necessity. CONSENT could therefore be the first self-correction method able to be applied to such ultra-long reads on a greater scale.

CONSENT’s assembly polishing feature also offers promising results. In particular, our experiment on a full human dataset shows that assembling the raw reads and then polishing the assembly allows to greatly reduce the computational costs, but also provides better results than assembling the corrected reads. This conclusion raises the question of the interest of long-read error correction in assembly projects. Moreover, as the processes of long read correction and assembly polishing are not much different from one another, one can also wonder why more error correction tools do not offer such a feature. It indeed seems to be affordable at the expense of minimal additional work, while providing satisfying results. We believe that CONSENT could open the doors to more error correction tools offering such a feature in the future. Finally, it would also be interesting to evaluate already published correction tools on their ability to polish assemblies, at the expense of minimal modifications to their workflows.

The segmentation strategy introduced in CONSENT also shows that actual multiple sequence alignments techniques are applicable to long, noisy sequences. In addition to being useful for error correction, this could also be applied to various other problems. For instance, it could be used during the consensus steps of assembly tools, for haplotyping, and for quantification problems. The literature about multiple sequence alignment is vast, but lacks application on noisy sequences. We believe that CONSENT could be a first work in that direction.

## Acknowledgements

Part of this work was performed using computing resources of CRIANN (Normandy, France).

