## Supplementary Material for "CONSENT: Scalable long read self-correction and assembly polishing with multiple sequence alignment"

Pierre Morisse[1,*], Camille Marchet[2], Antoine Limasset[2], Thierry Lecroq[3] and Arnaud Lefebvre[3]

[1]Normandie Université, UNIROUEN, INSA Rouen, LITIS, 76000 Rouen, France
[2]Univ. Lille, CNRS, UMR 9189 - CRIStAL
[3]Normandie Univ, UNIROUEN, LITIS, 76000 Rouen, France

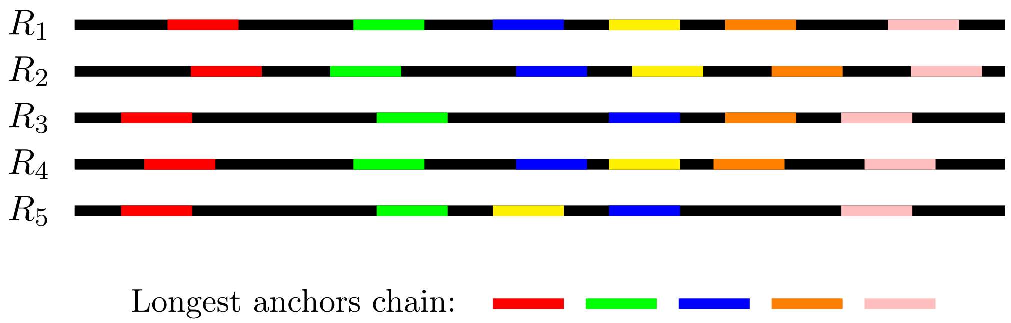


Figure S1: **Computation of the longest anchors chain for a set of sequences.** Here, we compute this chain with the second property set to $T = 3$, to simplify the figure. The longest chain computed here is thus $S =$ (red, green, blue, orange, pink). If we only considered the four first sequences, the chain $S' =$ (red, green, blue, yellow, orange, pink), which is long than $S$ could be selected. Indeed, although $R_3$ does not contain the yellow anchor, $S'$ follows the two properties in sequences $R_1$, $R_2$, and $R_4$. However, in $R_5$, the yellow anchor appears before the blue anchors, and thus invalidates the first property. $S'$ thus cannot be selected as the longest chain for the set of sequences $R_1$-$R_5$, and $S$ is chosen instead. Moreover, $R_5$ does not contain the whole set of anchors that appear in the longest anchor chain, $S$ (orange anchor is missing). As a result, $S_5$ will be filtered out during the MSA and consensus computation step.

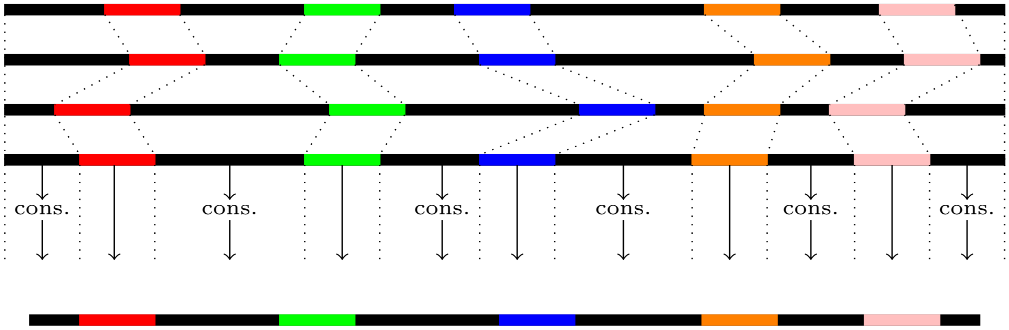


Figure S2: **Segmentation strategy for consensus computation of a window.** Anchors defining the longest anchors chain are used in order to segment the MSA and consensus computation. Local, independent consensus are thus computed on subsequenes delimited by these anchors. These local consensus, along with the anchors, are then concatenated, in order to rebuild to global consensus, at the scale of the original sequences length.

| Dataset | Number of reads | Average length (bp) | Error rate (%) | Coverage | Accession |
|---|---|---|---|---|---|
| **Simulated PacBio data** | | | | | |
| *E. coli 30x* | 16,959 | 8,235 | 12.29 | 30x | N/A |
| *E. coli 60x* | 33,918 | 8,211 | 12.28 | 60x | N/A |
| *S. cerevisiae 30x* | 45,198 | 8,216 | 12.28 | 30x | N/A |
| *S. cerevisiae 60x* | 90,397 | 8,204 | 12.29 | 60x | N/A |
| *C. elegans 30x* | 366,416 | 8,204 | 12.28 | 30x | N/A |
| *C. elegans 60x* | 732,832 | 8,220 | 12.28 | 60x | N/A |
| **Real ONT data** | | | | | |
| *D. melanogaster* | 1,327,569 | 6,828 | 14.55 | 63x | SRX3676783 |
| *H. sapiens*[1] (chr 1) | 1,075,867 | 6,744 | 17.60 | 29x | PRJEB23027 |
| *H. sapiens* | 15,243,243 | 7,411 | 17.26 | 35x | PRJEB23027 |

Table S1: Description of the long reads datasets used in our experiments.
[1] Only reads from chromosome 1 were used.

| Reference organism | Strain | Reference sequence | Size (Mbp) |
|---|---|---|---|
| *E. coli* | K-12 substr. MG1655 | NC_000913 | 4.6 |
| *S. cerevisiae* | W303 | scf7180000000{084-13} | 12.2 |
| *C. elegans* | Bristol N2 | GCA_000002985.3 | 100 |
| *D. melanogaster* | BDGP Release 6 | ISO1 MT/dm6 | 144 |
| *H. sapiens* (chr 1) | GRCh38 | NC_000001 | 249 |
| *H. sapiens* | GRCh38 | NC_000001 - NC_000024 | 3,257 |

Table S2: Description of the reference sequences used in our experiments.
[1] Only chromosome 1 was used.

| Dataset | Corrector | Throughput (Mbp) | Average length (bp) | Error rate (%) | Recall (%) | Precision (%) | Runtime | Memory (MB) |
|---|---|---|---|---|---|---|---|---|
| *E. coli* 30x | Original | 140 | 8,235 | 12.2862 | _ | _ | N/A | N/A |
| | LoRMA | 2 | 201 | 1.1962 | 99.9209 | 98.8208 | 10 min | 32,155 |
| *E. coli* 60x | Original | 279 | 8,211 | 12.2788 | _ | _ | N/A | N/A |
| | LoRMA | 171 | 886 | 0.1285 | 99.9865 | 99.8743 | 1 h 39 min | 31,682 |
| *S. cerevisiae* 30x | Original | 371 | 8,216 | 12.283 | _ | _ | N/A | N/A |
| | LoRMA | 14 | 248 | 2.1640 | 99.8351 | 97.8564 | 46 min | 31,899 |
| *S. cerevisiae* 60x | Original | 742 | 8,204 | 12.2886 | _ | _ | N/A | N/A |
| | LoRMA | 443 | 856 | 0.2225 | 99.9785 | 99.7812 | 5 h 25 min | 31,828 |
| *C. elegans* 30x | Original | 3,006 | 8,204 | 12.2806 | _ | _ N/A | N/A | |
| | LoRMA | 33 | 215 | 3.6960 | 99.7269 | 96.3449 | 8 h 19 min | 31,827 |
| *C. elegans* 30x | Original | 6,024 | 8,202 | 12.2825 | _ | _ | N/A | N/A |
| | LoRMA | 781 | 337 | 0.6446 | 99.9547 | 99.3649 | 31 h 04 min | 32,104 |

Table S3: Metrics output by ELECTOR on the simulated PacBio datasets, for the error correction with LoRMA. Runtime and memory consumption are reported for the whole correction pipeline. To underline the aggressive splitting of the reads performed by LoRMA, we also report the average length of the reads, in addition to ELECTOR's metrics.

On the 30x datasets, LoRMA performed worse than all the other methods, displaying throughputs 4x to 11x lower. This can be explained by the fact that it requires deep long-read coverage, as it is the only method purely relying on a $k$-mer strategy, thus fully avoiding overlaps computation. Indeed, LoRMA's throughput was much higher, and comparable to that of other methods, on the 60x datasets. However, it produced corrected reads displaying extremely short average length, barely reaching more than 800 bp on the 60x coverage datasets, and reaching less than 300 bp on the 30x coverage datasets. As for memory consumption, LoRMA consumed 32 GB, the maximum available amount of memory, on all the datasets.

| Dataset | Corrector | Number of reads | Throughput (Mbp) | N50 (bp) | Aligned reads (%) | Alignment identity (%) | Genome coverage (%) | Runtime | Memory (MB) |
|---|---|---|---|---|---|---|---|---|---|
| *D. melanogaster* | Original | 1,327,569 | 9,064 | 11,853 | 85.52 | 85.43 | 98.47 | N/A | N/A |
| | LoRMA | 1,125,279 | 6,386 | 669 | 97.05 | 98.47 | 94.76 | 23 h 51 min | 65,536 |
| *H. sapiens* (chr 1) | Original | 1,075,867 | 7,256 | 10,568 | 88.24 | 82.40 | 92.46 | N/A | N/A |
| | LoRMA | 737,198 | 1,247 | 186 | 96.50 | 97.83 | 28.62 | 13 h 07 min | 50,435 |

Table S4: Statistics of the real long reads, before and after correction with LoRMA. Runtime and memory consumption are reported for the whole correction pipeline.

In terms of alignment identity, LoRMA outperformed all the other tools. However, the N50 of the reads was extremely short, and did not even reach 700 bp, which is consistent the short read lengths observed in Supplementary Table S3. This confirms the fact that LoRMA tends to aggressively split the reads, and only manage to correct small portions of them. This small N50 thus explains the high alignment identities, by the fact that LoRMA tends not to correct complex regions of the reads. As a result, LoRMA also displayed the lowest genome coverage among all the tools, especially on the *H. sapiens* dataset, were the genome coverage was less than 30%. Assembly result for LoRMA corrected reads are thus not presented, since Miniasm could not manage to perform assembly with these reads.
